# OptoChaperone – A biohybrid tool for regulating protein condensates in cells and *in vitro*

**DOI:** 10.1101/2025.07.15.665009

**Authors:** Do Thanh Tuan, Motonori Matsusaki, Honoka Ota, Soichiro Kawagoe, Noriyoshi Isozumi, Hiroyuki Kumeta, Koichiro Ishimori, Eiichiro Mori, Tomohide Saio

## Abstract

Protein condensates formed via liquid–liquid phase separation (LLPS) are increasingly recognized as key players in diverse cellular processes, including those associated with disease. Despite extensive efforts to characterize their formation and function, tools that enable precise, reversible, and spatiotemporal control of LLPS remain limited. Here, we report OptoChaperone, a light-activatable molecular system designed to manipulate protein condensates both in vitro and in living cells. This biohybrid system leverages photoresponsive switching to control chaperone activity: blue light triggers the suppressive function, leading to the dissolution of protein condensates, whereas UV light deactivates the system, allowing condensate formation. We demonstrate the efficacy of OptoChaperone in regulating several disease-related protein condensates, such as fused in sarcoma, TAR DNA-binding protein 43, and heat shock factor 1. Importantly, the system exhibits reversible and robust control over droplet dynamics without requiring chemical additives or genetic modifications of the client proteins. Given the reversibility and efficiency of OptoChaperone in the manipulation of protein condensates, this tool offers a powerful platform for dissecting the roles of protein condensation in cellular physiology and pathology. This strategy also holds potential for broader applications in synthetic biology, biomolecular engineering, and therapeutic modulation of aberrant phase separation.

## Introduction

Protein aggregation is a hallmark of various neurodegenerative diseases^1–3^, including amyotrophic lateral sclerosis^4^, frontotemporal dementia^5^, Parkinson’s disease^6^, and Alzheimer’s disease^7,8^. Many disease-associated proteins, such as fused in sarcoma (FUS)^9^, TAR DNA-binding protein 43 (TDP-43)^10^, α-synuclein^11,12^, and tau^13^ undergo condensation, usually liquid–liquid phase separation (LLPS), as a prerequisite for pathological aggregation^14–16^. Protein condensation is also closely related to protein function as seen in stress response^17^, transcription regulation, RNA splicing^18^, and signal transduction^19^, where protein condensation assembles multiple components and increases the local concentration of the selected molecules. However, the functional significance of protein condensation remains unclear, and technologies elucidating the arbitrary control of protein condensates are required. Several factors, including pH, temperature, redox state, and salt concentration, can influence protein condensates^20–22^. However, these global environmental changes affect not only protein condensation but many other cellular events.

Optogenetic systems have offered a significant advancement in this field^23^. Several optogenetic systems have been developed to control protein condensate in cells, including optoDroplet^24^, PixELLs^25^, Corelet^26^, CasDrop^27^, iPOLYMER-L^28^, and LOVTRAP^29^. These systems typically function by fusing photoinducible oligomeric proteins with LLPS-potent proteins to enable light-induced droplet formation. Although these tools have advanced our understanding of protein condensates, they are primarily designed to induce droplet formation rather than disruption and, accordingly, often suffer from irreversible protein condensation. This limitation highlights the need for new tools to promote the disassembly of protein condensates.

Here, we propose a biohybrid optogenetic tool, OptoChaperone, that exploits a molecular chaperone to achieve reversible control of protein condensation and dissociation. Molecular chaperones have emerged as key factors in protein disassembly. For example, Hsp70 and Hsp27 are responsible for regulating TDP-43 and FUS condensates^30,31^. Karyopherin-β2, also known as a nuclear transport receptor, dissolves FUS condensates^32^. In this study, we designed OptoChaperone, utilizing a well-studied molecular chaperone trigger factor (TF) that recognizes unfolded proteins and intrinsically disordered proteins, including α-synuclein, to prevent the client proteins from self-assembly^33,34,35^. In OptoChaperone, the chaperone activity in the regulation of protein condensates is regulated by “lid protein,” composed of a protein labeled with an azobenzene derivative^36,37,38^. Photo-induced *trans/cis* isomerization of the azobenzene attached to the lid protein induces folding/unfolding of the lid protein, leading to exposure and coverage of the client-binding sites on the chaperone. Our data show that OptoChaperone reversibly controls protein condensation in cells and *in vitro*, as demonstrated using several disease-related protein condensates, including FUS^39^, TDP-43^40,41^, and heat shock factor 1 (Hsf1)^17,42,43^. Our results show that OptoChaperone provides an efficient strategy for studying protein condensates related to cellular processes and disease mechanisms.

## Results

### Design of OptoChaperone

OptoChaperone was constructed by fusing TF chaperone with the immunoglobulin-binding domain B1 of a streptococcal protein G (GB1) mutant labeled with a photo-switchable compound azobenzene derivative 3′-bis(sulfonato)-4,4′-bis(chloroacetamido)azobenzene (BSBCA)^36,44,45,46^, as a lid protein (**Figs 1a** and **b**). To immobilize BSBCA, the GB1 mutant was designed to have two exposed Cys residues (25 and 36) on the helix. The Cys residues were introduced at residue numbers *i* and *i* + 11, whose distance matches that of both ends of the BSBCA arms in the *trans* population^36,44^. In contrast, the *cis* population of BSBCA has a reduced arm distance^47^, leading to breakage of the helix^36,48^. Thus, the GB1 mutant labeled with BSBCA, azGB1, is expected to unfold with UV light, inducing the *cis* population of azobenzene. Given that TF binds unfolded proteins but not folded proteins^33^, it is expected to selectively bind *cis*-azGB1, resulting in buried client binding sites on TF. With *trans*-azGB1, the client-binding sites on the TF are expected to be exposed, creating an active state that binds client proteins and suppresses condensation. Accordingly, azGB1was used as a “lid” and fused to the TF domains: the TF^PPD-SBD^ consisting of peptidyl–prolyl isomerization domain (PPD: amino acid residues 150−246) and the substrate-binding domain (SBD: amino acid residues 113−149 and 247−432). Using the above design, the fusion protein azGB1-(Gly-Ser-linker)-TF^PPD-SBD^, hereafter referred to as OptoChaperone, was designed (**Fig 1c**). The details of the design and preparation are described in the Methods section.

**Fig 1.**
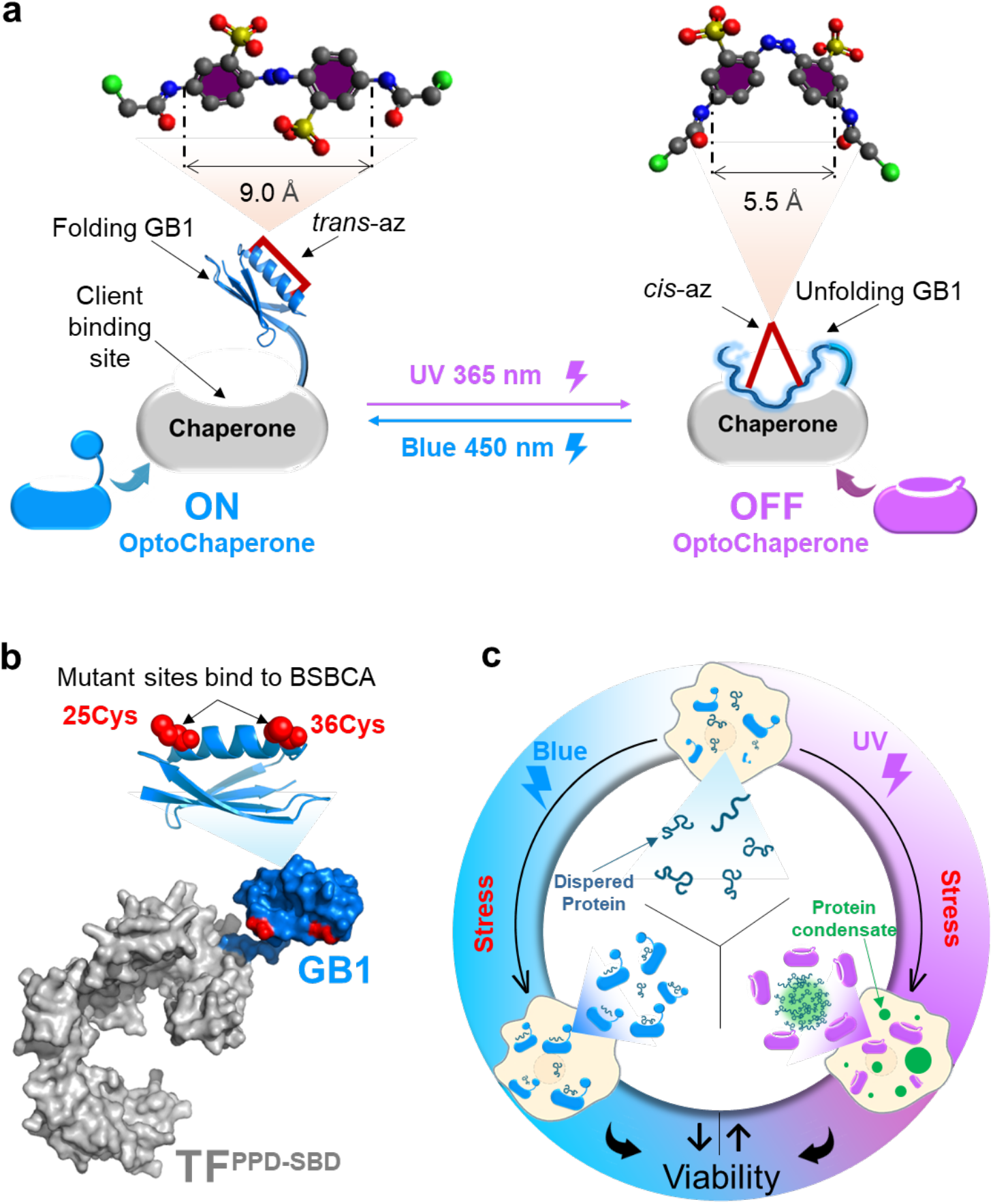
Schematic diagram of the new protein condensate light manipulation tool. **(a)** Structure of ON/OFF OptoChaperone and isomerization between different states by light irradiation. **(b)** Blueprint of a GB1-TF^PPD-SBD^ protein containing mutations for BSBCA introduction. Domain arrangement and structure of a full-length OptoChaperone include GB1-Solubilization tag with two Cys residue introduction for BSBCA modification; (GS)_5_-linker sequence; two domains of the trigger factor chaperone, peptidyl–prolyl cis/trans isomerase domain (PPD) residues numbered 150−246, and substrate-binding domain (SBD) residues numbered 113−149 and 247−432). **(c)** Conceptual model: OptoChaperone allows the reversible, light-dependent regulation of stress-responsive protein condensates in living cells, leading to modulation of stress response pathways and cell survival.

### Photo-response of OptoChaperone

The conformational state of the BSBCA moiety in the lid azGB1 after light irradiation was evaluated using UV-vis spectroscopy using the formula described in the Methods section (**Figs 2a** and **b**). The data showed that UV light increased *cis* conformation to ∼60%, whereas blue light induced *trans* conformation to ∼75% **(Fig 2b)**. These results indicated that the *cis*/*trans* populations of BSBCA could be induced using UV/blue light irradiation. Notably, the *cis*/*trans* population changes were repeatable, with no significant deterioration. The structure of the GB1 moiety of azGB1 after light irradiation was evaluated using nuclear magnetic resonance (NMR). The ^1^H-^15^N heteronuclear single quantum coherence (HSQC) spectra of ^15^N-labeled GB1 mutant attached to BSBCA (^15^N-azGB1) after blue light irradiation showed well-dispersed resonances, indicating that GB1 has a tertiary structure (**Fig 2c**). In contrast, the spectra after UV light irradiation showed reduced intensities, especially for the dispersed resonances, whereas the increased intensities for the resonances were located at approximately 8 ppm in the proton dimension. As the intense resonances with narrow dispersion around 8 ppm in the proton dimension are characteristic of unfolded proteins, the NMR data suggested that blue and UV lights induced the folded and unfolded states of azGB1, respectively. Thus, we concluded that light-induced *cis*/*trans* isomerization of BSBCA successfully modified the structure of GB1 attached to BSBCA.

**Fig 2.**
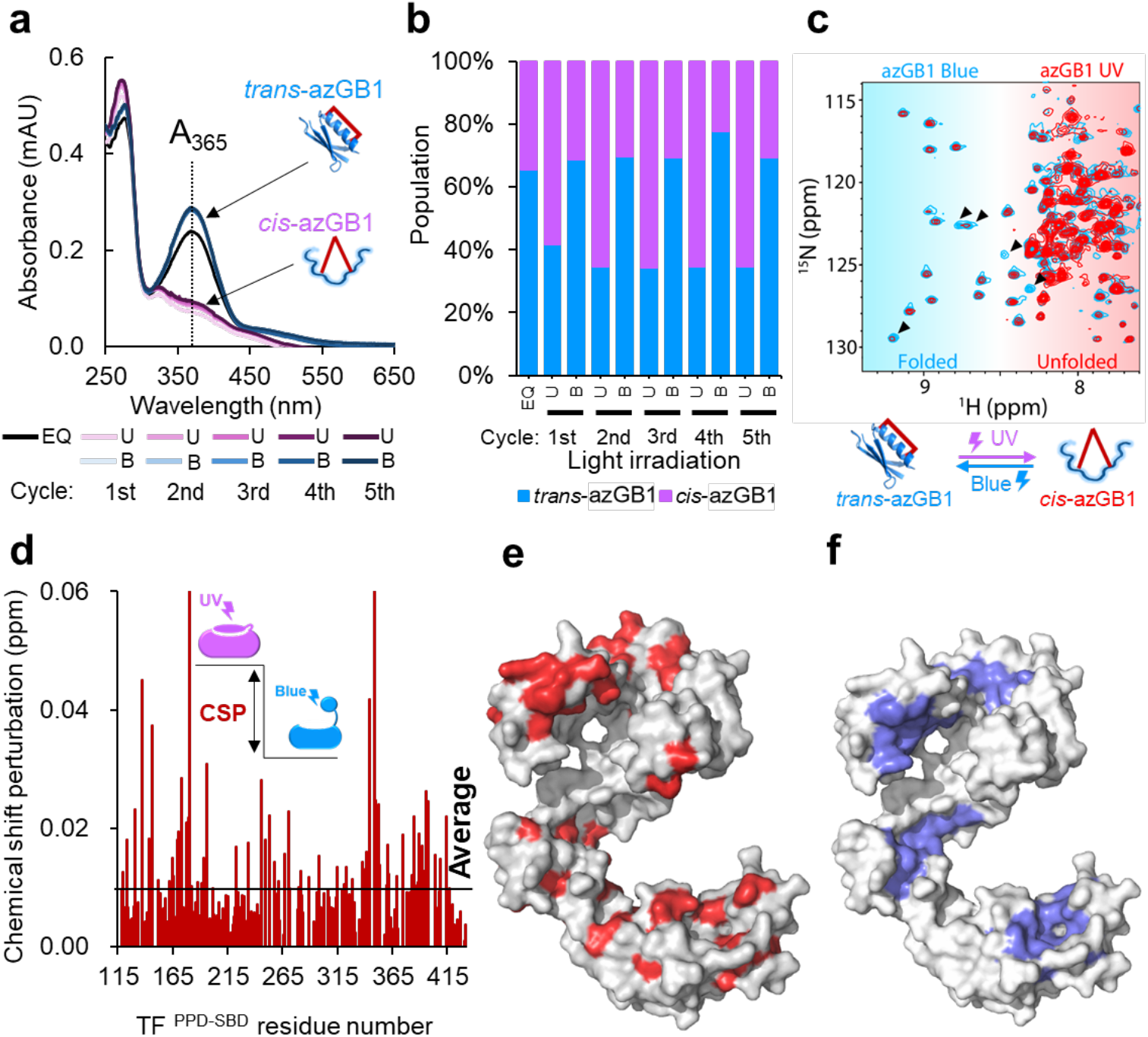
Investigation of properties of lid protein and OptoChaperone. **(a)** Ultra-visible (UV-vis) spectra of azGB1 measured after alternate irradiation using blue light and UV light for 5 min. **(b)** Comparison between the ratios of *trans*-azGB1 and *cis*-azGB1 after exposure to light. **(c)** Superposition of the [^1^H-^15^N] HSQC spectra of *trans-*azGB1 (blue) and *cis-*azGB1 (red). The blue signal is exposed to blue light, and the red signal is exposed to UV light. The arrowheads represent GB1-derived signals where significant changes in signal intensity were observed. **(d)** Chemical shift perturbation (CSP) of amide moieties between *cis*-azGB1-TF^PPD-SBD^ and *trans*-azGB1-TF^PPD-SBD^. The horizontal axis represents amino acid residues in OptoChaperone, and the vertical axis indicates the CSP. **(e)** Solvent accessible surface area (red) OptoChaperone between *cis*-azGB1-TF^PPD-SBD^ and *trans*-azGB1-TF^PPD-SBD^. **(f)** Mapping of the client-binding sites on the surface area (blue) of OptoChaperone. (EQ: Thermal Equilibrium, U: UV light irradiation, B: Blue light irradiation)

NMR data showed a photo-switchable interaction between azGB1 and TF^PPD-SBD^. Chemical shift differences were evaluated for resonances from TF^PPD-SBD^ (**Fig 2d**). Several specific residues, especially those from the client-binding sites,^33,34^ showed perturbations between the *trans* and *cis* states (**Figs 2e** and **f**). Additionally, the TF resonances from *cis*-OptoChaperone showed more significant perturbations from isolated TF^PPD-SBD^ compared to those from *trans*-OptoChaperone, indicating that *cis*-Optochaperone has a higher population of GB1 moieties bound to TF^PPD-SBD^ than *trans*-Optochaperone. These data suggest that photoswitching of the GB1 structure by BSBCA controls the interaction between GB1 and TF^PPD-SBD^, highlighting the ability of azGB1 to function as a lid for TF chaperones. Regarding chaperone activity, *trans*-OptoChaperone should serve as an ON state or active state, while *cis*-OptoChaperone should serve as an OFF state or inactive state.

### OptoChaperone regulates protein condensates in vitro

The role of OptoChaperone in regulating protein condensates was evaluated *in vitro*. We first attempted to demonstrate protein condensate regulation by microscopic observation of Hsf1 protein condensate induced in the presence or absence of the ON/OFF state OptoChaperone or unmodified chaperone, TF^PPD-SBD^ (**Fig 3a**). With ON-state OptoChaperone, droplet formation was suppressed, and only small droplets were observed even after 60 min, indicating that OptoChaperone successfully suppressed Hsf1 assembly (**Fig 3a**). In contrast, OFF-OptoChaperone exhibited negligible suppression and allowed droplet formation. This result strongly supports the design of the OptoChaperone: UV light induces OFF-state OptoChaperone, losing its ability to bind Hsf1 and triggering Hsf1 condensation. Conversely, blue light induces ON state OptoChaperone, which captures Hsf1 and prevents Hsf1 self-assembly. The same trend was observed in the turbidity assays (**Fig 3b**). In the presence of OptoChaperone after blue light (ON state), the turbidity was maintained at ∼0.2, whereas after UV light irradiation, inducing the OFF state, the turbidity increased to ∼0.5, indicating that the ON state OptoChaperone successfully suppressed Hsf1 assembly, whereas the OFF state OptoChaperone allowed assembly. Furthermore, the data showed that the formation and suppression of Hsf1 droplets were reversible (**Fig 3b**). Turbidity decreased after 5-min irradiation with blue light, indicating the suppression of Hsf1 droplets. In contrast, turbidity increased after 5 min of UV light irradiation, indicating a reduction in OptoChaperone activity. These reversible effects were observed for up to three cycles, demonstrating the reversible switching of chaperone activity.

**Fig 3.**
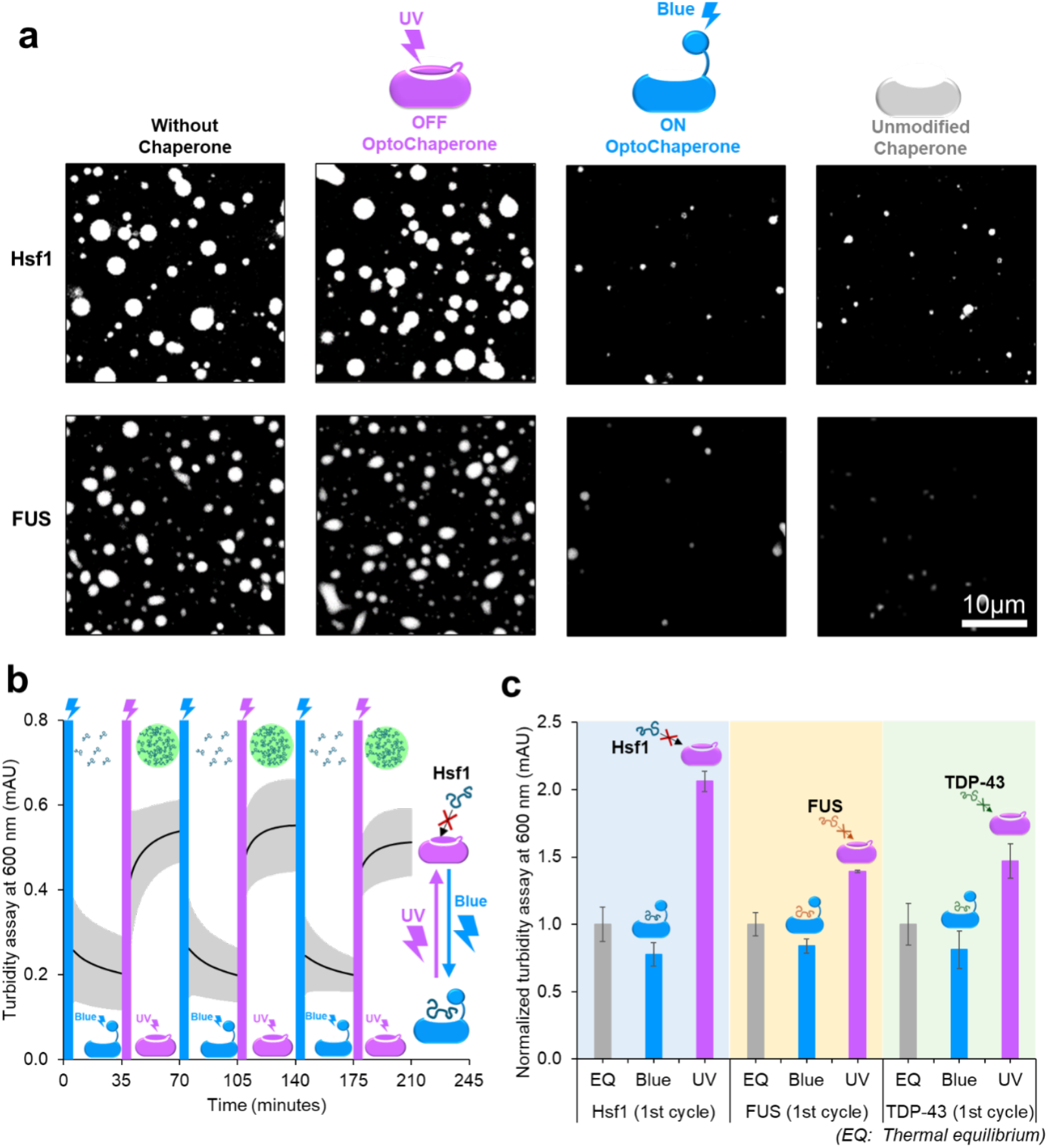
OptoChaperone enables reversible *in vitro* control of protein condensate formation. **(a)** Confocal microscopy images of heat shock factor 1 (Hsf1) and fused in sarcoma (FUS) droplets at 60 min. For OFF and ON OptoChaperone conditions, samples were irradiated for 5 min with UV (OFF) or blue (ON) light. Samples contain purified protein (Hsf1 or FUS with 10% fluorescent-labeled protein) with or without equivalent concentrations of OFF OptoChaperone, ON OptoChaperone, or unmodified chaperone. Droplet formation was triggered using crowding agents: 10% w/v Ficoll (Hsf1) and 4% w/v PEG8000 (FUS). Scale bar: 10 μm. **(b)** Time-course turbidity measurements (600 nm) of Hsf1 solutions during three cycles of alternating UV and blue light exposure (5 min each). After 30 min of adding a crowding agent or light irradiation, the next light irradiation was performed for 5 min, and the turbidity was measured again. Error bars: standard deviation (n = 3). **(c)** Normalized turbidity of Hsf1, FUS, and TAR DNA-binding protein 43 (TDP-43) solutions at 30 min after initial UV or blue light exposure (1st cycle). Values normalized to initial turbidity before light exposure, set as 1.0 (n = 3).

Photoswitching of protein condensation was also demonstrated using FUS and TDP-43. Microscopic observations showed that OptoChaperone was also capable of controlling FUS droplets, with significant differences in terms of the size and number of FUS droplets between the ON and OFF states (**Fig 3a**). The turbidity assay corroborated the microscopic observations, showing significant changes in turbidity after light irradiation (**Fig 3c**). A turbidity assay was also performed for TDP-43, showing that OptoChaperone is effective in regulating the TDP-43 condensate (**Fig 3c**).

Collectively, *in vitro* assays demonstrated that OptoChaperone is effective for Hsf1, FUS, and TDP-43 proteins, suppressing droplet formation after blue light, which triggers the ON state of OptoChaperone and allows droplet formation after UV light, which triggers the OFF state of OptoChaperone. Nevertheless, the effects on these proteins could differ owing to variations in the substrate recognition mechanisms of TF. Taking advantage of the broad client recognition properties of TF, OptoChaperone, consisting of TF^PPD-SBD^, can handle many different disordered proteins.

### OptoChaperone regulates protein condensates in cells

Subsequently, to evaluate whether OptoChaperone regulates the protein condensate of Hsf1 in cells, OptoChaperone was introduced into HeLa cells, and Hsf1 condensation after blue/UV light irradiation was observed using confocal microscopy (**Fig 4a**). After electroporation followed by a 3-h incubation period at 37 °C, the cells were exposed to blue and UV light for 5 min and subsequently treated under heat at 43 °C for 1 h. Endogenous Hsf1 was observed using immunostaining. Confocal images of the cells after heat treatment showed that a larger number and size of Hsf1 foci were observed for cells containing OFF-state OptoChaperone than for those containing ON-state OptoChaperone (**Fig 4b**). Evaluation of the foci area corroborated this observation, showing that cells containing OFF-state chaperones induced by UV irradiation possessed a larger area of Hsf1 foci than that in those containing ON-state chaperones induced by blue light (**Fig 4c**). These observations were highly consistent with *in vitro* results and highlighted the ability of OptoChaperone to function under cellular conditions.

**Fig 4.**
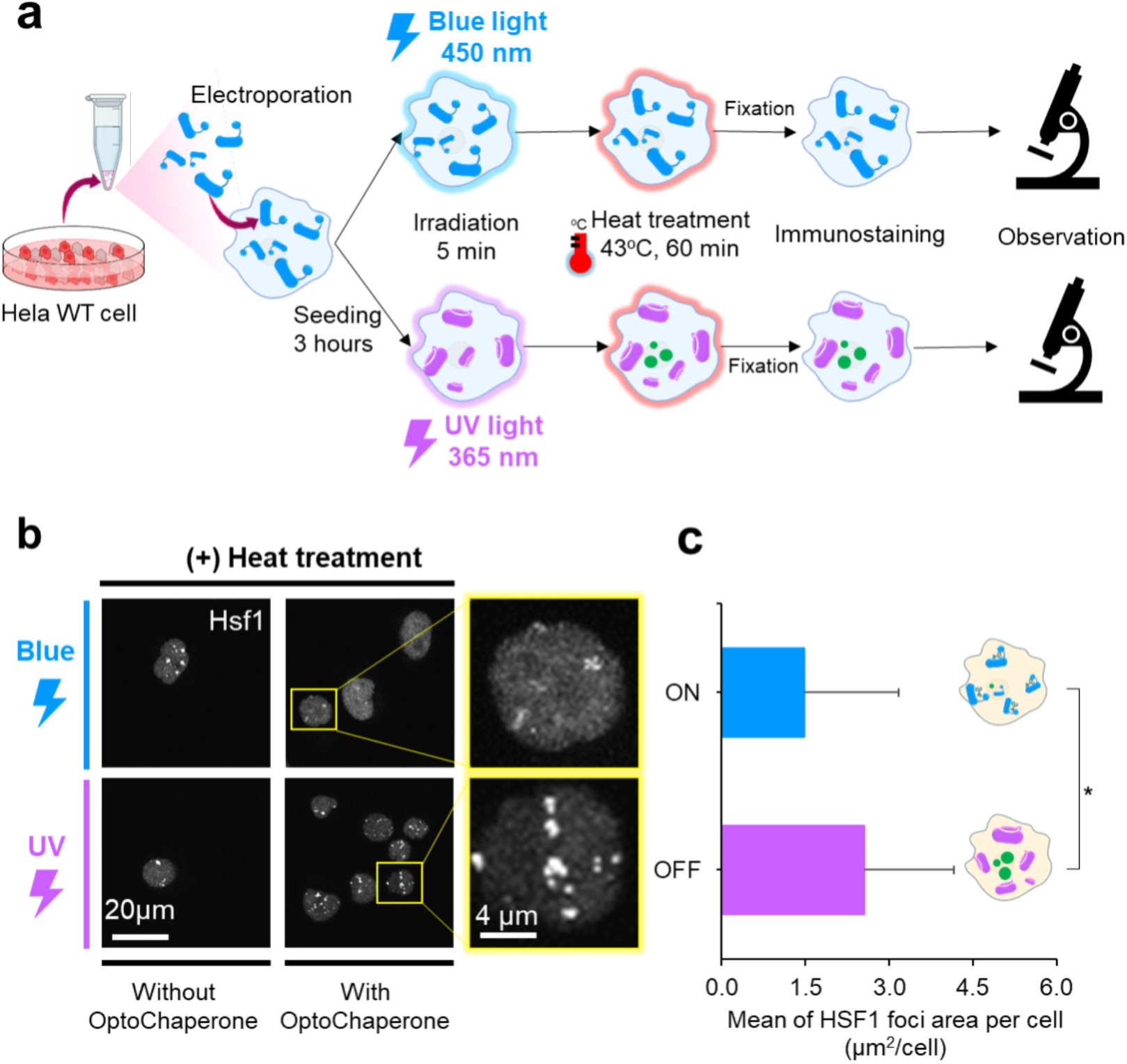
OptoChaperone-mediated control of protein condensates in cells. **(a)** Schematic of the experimental workflow for observing fixed intracellular foci formation. HeLa cells were electroporated with OptoChaperone, incubated for 3 h at 37 °C, and then exposed to either 5 min of UV or blue light. All groups were subsequently subjected to 60-min heat shock, fixed, immunostained, and observed. **(b)** Representative confocal images of endogenous Hsf1 foci in OptoChaperone-introduced cells (Scale bar 20 μm). Enlarged images of the yellow boxed areas are shown (Scale bar 4 μm). Hsf1 was detected using an anti-Hsf1 antibody. **(c)** Quantitative analysis of Hsf1 foci formation per HeLa cell containing OptoChaperone after light irradiation and heat shock (43 °C, 60 min). Error bars: standard deviation (ON: n = 20, OFF: n = 27). Statistical significance was determined by unpaired, two-tailed Student’s *t*-test (*p < 0.05).

### Photo-switching of OptoChaperone determines the fate of cells under stressed conditions

Building on the regulatory role of OptoChaperone in cells (**Fig 4**), we evaluated the effect of OptoChaperone activity in cells. Given that the stress response induces the formation of several protein condensates, including nuclear stress bodies containing Hsf1 and stress granules containing FUS and TDP43, the photoswitching of OptoChaperone was expected to affect the stress response system. Therefore, we evaluated the cell viability after heat treatment in the absence and presence of ON/OFF-OptoChaperone (**Fig 5)**. The cell viability of the OFF-OptoChaperone group was comparable to that of the non-OptoChaperone group but significantly higher than that of the ON-OptoChaperone group. The reduced cell viability with ON-OptoChaperone can be attributed to the suppression of the stress-responsive protein condensates as represented by Hsf1 condensates (**Fig 4**), confirming the critical function of these protein condensates for cells to survive under stress conditions.^49^ Thus, the data showed that OptoChaperone not only controls protein condensates but also controls cell fate under stressed conditions, demonstrating its effectiveness in studying the functional importance of protein condensates in the cell.

**Fig 5.**
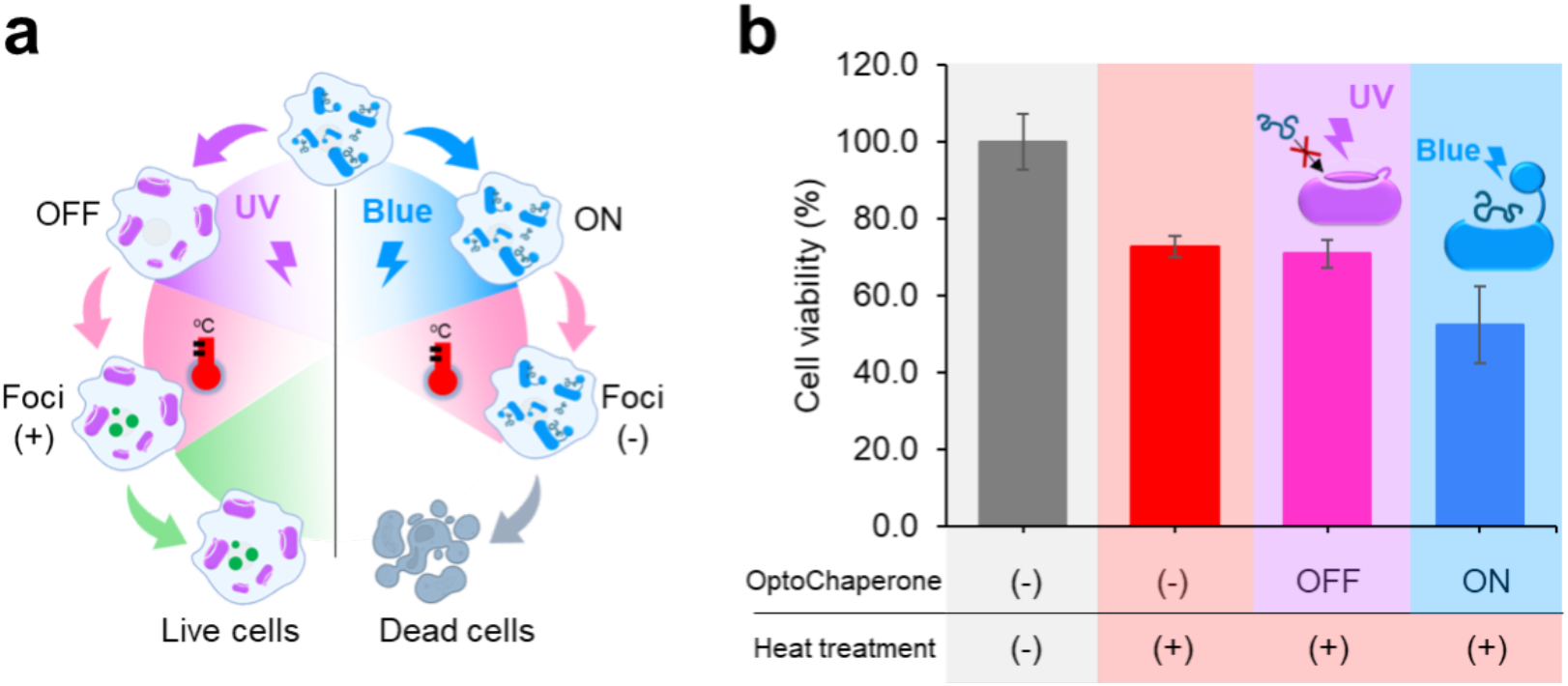
Effect of OptoChaperone on cell viability under heat treatment. **(a)** Schematic of cell viability assay. HeLa cells electroporated with OptoChaperone were seeded at 16,000 cells/well in collagen type I-coated 96-well plates. After adherence, cells were irradiated for 5 min with UV (OFF) or blue (ON) light to switch the state of OptoChaperone before heat treatment. Cells were then subjected to 120 min of heat treatment at 43 °C, followed by a 44-h incubation at 37 °C. Cell viability was measured using a water-soluble tetrazolium 8 (WST-8) assay after 4 h of incubation with the reagent. **(b)** Quantitative analysis of cell viability after light irradiation and heat treatment. Viability values were normalized to those of HeLa cells without OptoChaperone and without heat treatment (set as 100%, n = 3). Error bars indicate standard deviation.

## Discussion

Unlike traditional methods, which rely on altering environmental conditions such as temperature, pH, ionic strength, and molecular crowding to induce protein condensation^20–22^, OptoChaperone leverages light to exert spatiotemporal control over protein condensates, allowing researchers to induce or reduce protein condensates without disturbing the cellular environment. One of the key strengths of the OptoChaperone is its reversibility. While previous optogenetic tools driving protein assembly often result in irreversible protein condensation^50^, OptoChaperone accelerates the dissociation of the protein condensate, which ensures reversible regulation. Given its unique concept and novel intervention points, OptoChaperone can be combined with other optogenetic tools that induce protein assembly using light. This extends the utility and potential application of optogenetic tools to study protein condensate dynamics and function in cells and *in vitro*. OptoChaperone offers advantages in controlling protein condensation through light, but the *cis/trans* switching of BSBCA requires UV light, which may raise concerns about potential cellular damage^51,52^. As evidenced by the results in **Fig 5**, the impact of UV and blue light on cells was minimal; however, this issue will be adequately addressed in future studies by using new azobenzene compounds responding to visible or near-infrared light^53^ that are less harmful to the cell.

Although we developed a non-selective OptoChaperone with a promiscuous chaperone TF, selective light irradiation to specific regions of cells would achieve special selectivity. Additionally, the same strategy can be applied to other chaperones, including those working for specific targets, by fusing a lid protein labeled with BSBCA because many chaperones share the ability to recognize unfolded hydrophobic regions of the proteins. This opens up new applications for this tool. The application of OptoChaperone is not limited to protein condensate studies but is also beneficial in the functional analysis of a chaperone in the cell. Given the increasing interest in chaperones related to diseases involving protein aggregation, OptoChaperone will be an important tool in broad research fields, including neurodegenerative diseases. OptoChaperone provides a platform for developing therapeutic strategies targeting pathological protein condensates, paving the way for new research directions and clinical applications.

## Methods

### Plasmids construction, protein expression, and purification

GB1 was expressed in the pET21d(+) vector. GB1-His_6_-TEV-GB1-(GS)_5_-TF^PPD-SBD^: GB1-(GS)_5_-TF^PPD-SBD^ was cloned into the pET21d(+) vector and fused with a GB1-His_6_ tag at the N-terminal TEV protease cleavage site. GB1-His_6_-Hsf1: A codon-optimized recombinant plasmid encoding Hsf1 was synthesized by Kawagoe et al.^43^ Hsf1 constructs were cloned into the pET21b vector (Cat. No. 69741-3CN, Novagen) and fused to a GB1-His_6_ tag at the HRV3C N-terminal PreScission protease cleavage site. MBP-TEV-FUS: FUS proteins were expressed in the pMAL-TEV vector using the MBP fusion constructs. MBP-TEV-TDP-43: TDP-43 proteins were expressed using MBP fusion constructs in the pMAL-TEV vector. All the recombinant proteins were expressed individually in BL21(DE3) cells. All the expression constructs were transfected into BL21(DE3) cells. Cells were grown in Luria−Bertani medium at 37 °C in the presence of ampicillin (50 μg mL^-1^). Subsequently, protein expression was induced by adding 0.5 mM isopropyl-β-D-1-thiogalactopyranoside (IPTG) at OD_600_ ∼0.6, followed by 12−16 h of incubation at 18 °C. The cells were harvested at OD_600_ ∼3.0, resuspended in lysis buffer containing 50 mM Tris-HCl (pH 8.0) and 500 mM NaCl, and stored at -80 °C. His_6_-GB1 and GB1-His_6_-TEV-GB1-(GS)_5_-TF^PPD-SBD^ were transfected into BL21(DE3) cells for recombinant protein expression. The cells were disrupted using a sonicator and centrifuged at 18000 rpm for 30 min. The supernatant containing the target protein was purified using a Ni-NTA Sepharose column (Cat. no. 30230; QIAGEN, Germany). After washing (using 50 mM Tris pH 8.0, 500 mM NaCl, 50 mM Tris pH 8.0, 150 mM NaCl, and 15 mM imidazole), the proteins were eluted with an Elution Buffer (50 mM Tris pH 8.0, 150 mM NaCl, and 400 mM imidazole). TEV protease was used to remove the purification tag, followed by overnight dialysis (50 mM Tris HCl, 150 mM NaCl, 4 mM 2-ME, pH 8.0) at 4 °C. The protein was further purified using a Ni-NTA resin (Cat. no. 30230; QIAGEN, Germany). Size-exclusion chromatography was performed using a HiLoad 26/600 Superdex 200 pg column (Cat. No. 10250637, Cytiva, Sweden; Buffer 50 mM Tris-HCl, 100 mM NaCl, pH 8.5). The purified protein was collected and stored at - 80 °C. The expression and purification of Hsf1^43^, FUS^32^, and TDP-43 proteins^41^ were performed according to a previous study.

### Protein isotope labeling for NMR studies

^15^N-isotopically labeled GB1 for NMR studies was prepared by growing the cells in a minimal (M9) medium containing ^15^NH_4_Cl (1 g liter^−1^). Protein expression was induced by adding 0.5 mM IPTG at OD_600_ ∼0.6, followed by 12−16 h of incubation at 18 °C. The cells were harvested at OD_600_ ∼2.0, resuspended in lysis buffer containing 50 mM Tris-HCl (pH 8.0) and 500 mM NaCl, and stored at -80 °C. For the isotope labeling of TF^PPD-SBD^ and GB1-(GS)_5_-TF^PPD-SBD^, the M9 medium containing ^15^NH_4_Cl and [^2^H,^12^C]-glucose in 99.9% D_2_O was used. For the production of U-[^2^H], Ile-δ1-[^13^CH_3_], Val,Leu-[^13^CH_3_,^12^CD_3_], Met-[^13^CH_3_], and Ala- [^13^CH_3_] TF^PPD-SBD^ and GB1-(GS)_5_-TF^PPD-SBD^, alpha-ketobutyric acid [methyl-^13^CH_3_] (50 mg liter^−1^), alpha-ketoisovaleric acid [monothyl-^13^CH_3_,^12^CD_3_] (85 mg liter^−1^), [^13^CH_3_]-Met (50 mg liter^−1^), and [D_2_, ^13^CH_3_]-Ala (50 mg liter^−1^) were added to the culture 30 min before adding IPTG. Protein expression was induced by adding 0.5 mM IPTG at OD_600_ ∼0.6, followed by 12−16 h of incubation at 18 °C. The cells were harvested at OD_600_ ∼2.0, resuspended in lysis buffer containing 50 mM Tris-HCl (pH 8.0) and 500 mM NaCl, and stored at -80 °C. The samples were purified, and BSBCA labeling was performed using the same procedure as that used for culturing in a Luria−Bertani medium.

### Introduction of BSBCA into GB1 or GB1-(GS)_5_-TF^PPD-SBD^

In a nitrogen atmosphere, a glove box was used to introduce BSBCA into a GB1 or GB1-(GS)_5_-TF^PPD-SBD^ solution. TCEP was then added, and the mixture was stirred at room temperature for 30 min. BSBCA was then added and stirred at 25 °C for 6 h. According to the previous study^36^, the *cis* and *trans* populations were determined from absorbance values. The *trans* concentration was calculated using Equation 1, assuming a negligible *cis* absorbance at 365 nm. The *cis* concentration was obtained using Equation 2, while the BSBCA in *trans-* and *cis-* populations were determined using Equations 3 and 4, respectively.

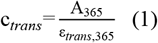

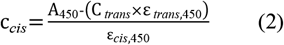

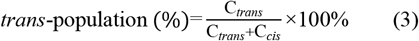

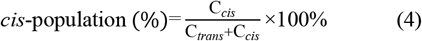

where C_*trans*_: Concentration of *trans*-population; C_*cis*_: Concentration of *cis*-population, A_365_: Absorbance of azGB1 at 365 nm; A_450_: Absorbance of azGB1 at 450 nm; ε_*trans*,365_: Extinction coefficient of *trans* at 365 nm (=24000 M^-1^. cm^-1^); ε_*trans*,450_: Extinction coefficient of *trans*-population at 450 nm (=2100 M^-1^. cm^-1^); ε_*cis*,450_: Extinction coefficient of *cis*-population at 450 nm (=2900 M^-1^. cm^-1^).

### Matrix-assisted laser desorption/ionization-time of flight mass spectrometry (MALDI-TOF-MS)

The samples were mixed with a saturated solution of sinapinic acid in 50% acetonitrile and 0.1% trifluoroacetic acid at a 1:1 ratio (v/v). A 1.0 μL aliquot of the mixture was spotted onto a MALDI target plate and air-dried at room temperature. Mass spectrometry analysis was performed using an Autoflex Speed (Bruker) with a Smartbeam II laser at the Center for Technical Support, Graduate School of Technology, Industrial and Social Sciences, Tokushima University (Tokushima, Japan). Ionization was performed in positive linear mode with an acceleration voltage. Mass spectra were acquired over an m/z range of 5–20 kDa using Bruker Daltonics flex analysis. Mass spectra were analyzed using Bruker Daltonics Flex Analysis to identify peaks corresponding to biomolecules of interest.

### NMR experiments

NMR samples were prepared in 20 mM KPi (pH 7.0), 100 mM KCl, 4 mM βME, 0.5 mM EDTA, 0.05% NaN_3_, and 7% D_2_O. Protein concentrations were 0.02 mM azGB1, 0.3 mM TF^PPD-SBD^, and 0.5 mM azGB1-TF^PPD-^ _SBD_. NMR experiments were performed using a Bruker Avance Neo 800 MHz NMR spectrometer with cryogenic probes and a Bruker Avance III 500 MHz NMR spectrometer. Following initial NMR acquisition, UV/blue light irradiation was applied for 5 min. Spectra were processed using the NMRPipe program,^54^ and data analysis was performed using Olivia (Fermi Pharm Hokudai Acc Jp/Olivia).

### Microscopic observation

For Hsf1 droplet formation, a solution containing 18 μM Hsf1, 2 μM GFP-Hsf1, 20 μM OptoChaperone, and 10 mM DTT was prepared. The solution was incubated at room temperature for 10 min before adding Ficoll400 to induce droplet formation. For MBP-FUS droplets, a solution of 4.5 μM FUS, 0.5 μM CF488A-FUS, 10 μM OptoChaperone, and 10 mM dithiothreitol (DTT) was prepared, followed by incubation for 10 min. PEG8000 was added to trigger droplet formation. Microscopic observations were conducted immediately using a laser scanning microscope equipped with a 40× objective lens. Light irradiation (5 min) for OptoChaperone activation was performed under the same conditions as the absorption spectrum measurements using a light guide positioned between an aluminum foil-shaded observation dish and the microscope. The radius and area occupancy ratio of the droplets in the image were measured using the ImageJ software, where the area-occupancy ratio is the proportion of the area occupied by each droplet within the field of the microscope.

### Turbidity assay

Turbidity assays were performed using a V-730 spectrophotometer at 25 °C with an initial blank measurement in the corresponding purification buffer after 5 min of light irradiation. Hsf1 (20 μM), FUS (5 μM), and TDP-43 (10 μM) solutions were prepared in their respective purification buffers with OptoChaperone (equal molar ratio) and DTT (10 mM) and incubated at room temperature for 30 min. Droplet formation was induced by adding Ficoll (10% w/v) to Hsf1 or PEG8000 (4% w/v for FUS and 8% w/v for TDP-43), followed by pipette mixing. The solutions (400 μL) were transferred to a quartz microcell, and turbidity (600 nm) was recorded every 30 s for 30 min. After the measurement, the samples were irradiated with light, shielded with aluminum foil, and re-measured under identical conditions.

### Cell culture

HeLa cells were obtained from RIKEN BRC, which is participating in the National Bio-Resource Project of MEXT, Japan. The HeLa cells were cultured and maintained at 37 °C and 5% CO_2_ in Dulbecco’s Modified Eagle’s Medium (DMEM) (Cat. No. L4E6239; Nacalai Tesque) supplemented with 10% fetal bovine serum (FBS) (Cat. No. F7254; Sigma-Aldrich, Tokyo, Japan). Once the cells proliferated to cover the dish, they were passaged every 2–3 days. The medium was removed, and the cells were washed twice with phosphate-buffered saline (PBS). Trypsin (5% in PBS) was added to detach cells, followed by incubation for 5 min. Trypsin was neutralized with 8 mL of DMEM, and the suspension was centrifuged (100 rpm for 3 min). The supernatant was removed, and the pellets were resuspended in 2 mL of DMEM. A 15 μL sample was used for counting in a hemocytometer, and cell density was calculated before reseeding in fresh DMEM. The cell suspension was adjusted so that the cell density per dish was 6×10^5^, and the cells were seeded.

### Cellular introduction of OptoChaperones using electroporation

The procedure involved adjusting the liquid volume to achieve a cell density of 1.0×10^6^ cells before seeding. The cells were transferred to a 1.5 mL tube, followed by centrifugation at 100 rpm for 3 min, and the supernatant was removed. They were then washed twice with 1 mL of electroporation buffer and mixed with 100 μL of 500 μM OptoChaperone solution (50% fluorescence labeling). The suspension was transferred to an electroporation cuvette, warmed to 37 °C, and subjected to electroporation using NEPA21 settings. The poring pulse was set at 100 V for 15 ms, and the transfer pulse was set at 20 V for 50 ms with alternating polarity. Finally, 200 μL of pre-warmed DMEM was added for recovery.

### Light irradiation

For UV exposure, a Xenon light source (Model: MAX-303, Asahi Spectra, Japan) was configured to emit monochromatic UV light isolated using a 365 nm bandpass filter at its maximum power output for 5 min. Subsequently, the light source was transitioned to emit blue light, filtered using a 450 nm bandpass filter, again at maximum power output, for an additional 5-min exposure period. The distance between the MAX-303 light source and sample was kept constant throughout both irradiation steps to ensure a consistent energy fluence.

### Cell viability assay

Wild-type HeLa cells (with OptoChaperone via electroporation) were seeded at a density of 16,000 cells/well in a 96-well collagen type I (Cat. No. 2962401; AGC Techno (Iwaki) Glass Co., Shizuoka, Japan) coated microplate. Each well contained 100 μL of DMEM. Cells were allowed to adhere for 2 h at 37 °C and 5% CO_2_. Following adherence, cells underwent either UV or blue light irradiation for 5 min, followed by heat treatment at 43 °C for 120 min. Subsequently, cells were incubated at 37°C and 5% CO_2_ for 44 h, with fresh DMEM introduced after 24 h. After the 44-h incubation, 100 μL of DMEM (pre-warmed to 37 °C) containing 10% WST-8 reagent (Cat. No. LD1023R; Kishida Chemical Co., Osaka, Japan) was added to each well. The plate was then incubated for an additional 4 h. The absorbance was measured at 450 nm using a microplate reader to assess cell viability.

## Acknowledgments

We appreciate Dr. Hitoki Nanaura, Dr. Honoka Kawamukai, and Dr. Haojie Zhu for their involvement in the initial experiments since the conception of this project. We also thank Ms. Eri Sakamoto and Ms. Aiko Iwano for their help with cell culture. This study used experimental resources provided by the Fujii Memorial Institute of Medical Sciences, Institute of Advanced Medical Sciences, and Tokushima University, Japan. This work was supported by the Medical Research Center Initiative for High Depth Omics, the Joint Usage and Joint Research Programs, and the Institute of Advanced Medical Sciences (IAMS) at Tokushima University. Some of the NMR experiments were performed at the Hokkaido University Advanced NMR facility, which is a member of the NMR Platform. This work was supported by funding from JSPS KAKENHI (JP24K18063 to S.K., JP20K15969, JP22H04847, and JP22K15278, and JP23KK0105 to M.M., JP19K06504, JP19H04945, JP20H03199, JP20KK0156, JP22H02560, JP22K18361, JP23H05470, JP23H01995, JP23K23824, and 25K02217 to T.S., and JP19H05769 to K.I.), MEXT Grant-in-Aid for Transformative Research Areas (B) (JP21H05094 and JP21H05093 to T.S.), AMED (JP22ek0109437, 25ek0109642 to T.S. and JP24wm0425004 to T.S. and E.M.), and JST FOREST Program (JPMJFR204W to T.S.). This work was supported by funding from JST SPRING, Grant Number JPMJSP2113 (D.T.T.) and the Home for Innovative Researchers and Academic Knowledge Users Driving Global Impact (HIRAKU)-Global Program, which is funded by MEXT’s “Strategic Professional Development Program for Young Researchers” (M.M.). This work was also partially supported by the Astellas Foundation for Research on Metabolic Disorders and the Uehara Memorial Foundation (M.M.) and the Takeda Science Foundation Grant, Sumitomo Foundation, Astellas Foundation for Research on Metabolic Disorders, Senri Life Science Foundation, Nakajima Foundation, Asahi Glass Foundation, Nakabayashi Trust for ALS Research, Kato Memorial Trust for Nambyo Research, Mochida Memorial Foundation for Medical and Pharmaceutical Research, The Canon Foundation, JKA Foundation, Naito Foundation, and Serika Fund (T.S).

## Author contributions

T. Saio and E. Mori conceptualized and planned the study. T. Saio, M. Matsusaki, and K. Ishimori supervised the project. T. Saio, M. Matsusaki, D. T. Tuan, H. Ota, and S. Kawagoe designed the experiments. D.T. Tuan, H. Ota, M. Matsusaki, and N. Isozumi performed biochemical experiments. S. Kawagoe and H. Kumeta performed the NMR experiments and analyzed the NMR data. D.T. Tuan, H. Ota, and M. Matsusaki analyzed the biochemical data and interpreted the data with T. Saio. D.T. Tuan, M. Matsusaki, and T. Saio wrote the manuscript. M. Matsusaki and T. Saio edited and revised the manuscript. All the authors have read and approved the final manuscript.

